# Backward masking in mice requires visual cortex

**DOI:** 10.1101/2021.09.26.461573

**Authors:** Samuel D Gale, Chelsea Strawder, Corbett Bennett, Stefan Mihalas, Christof Koch, Shawn R Olsen

## Abstract

Visual masking is used to infer the timescale of conscious perception in humans; yet the underlying circuit mechanisms are not understood. Here we describe a two-alternative choice backward masking task in mice and humans, in which the location of a briefly presented grating is effectively masked within a 50 ms window after stimulus onset. Importantly, human subjects reported reduced perceptual visibility during masking that closely corresponded with behavior deficits. In mice, optogenetic silencing of visual cortex reduces performance over a similar time course as masking. However, response rates and accuracy do not match masking, demonstrating that cortical silencing and masking are distinct phenomena. Spiking responses recorded in primary visual cortex (V1) are consistent with masked behavior when quantified over long, but not short, time windows, indicating masking involves further downstream processing. Behavioral performance can be quantitatively recapitulated by a dual accumulator model constrained by V1 activity. The model and the animal’s performance for the earliest decisions imply that the initial spike arriving from the periphery can trigger a correct response, but subsequent V1 spikes, evoked by the mask, degrade performance for longer decisions. To test the necessity of visual cortex for backward masking, we optogenetically silenced mask-evoked cortical activity which fully restored discrimination of target location. Together, these results demonstrate that mice, like humans, are susceptible to backward visual masking and that visual cortex has a crucial role in this process.

## Introduction

Visual masking has been used to probe the temporal dynamics of conscious perception and unconscious processing for well over a century (Bachmann and Francis, 2013; Breitmeyer and Ogmen, 2006; Dehaene, 2014). In backward masking, the visibility of a target is suppressed by a subsequent stimulus, the mask. This retroactive influence of the mask on the target can be accounted for by multiple mechanisms, including lateral inhibitory interactions between feedforward circuits in the visual pathway (Alpern, 1953; Battersby and Sturr, 1970; Breitmeyer and Ogmen, 2006; Macknik and Martinez-Conde, 2004a; Öǧmen, 1993), disruption of cortical feedback (Lamme et al., 2002; Ro et al., 2003), or long temporal integration of feedforward sensory signals causing the perception of target and mask to merge (Eriksen and Hoffman, 1963; Schultz and Eriksen, 1977; Thompson, 1966). The inter-ocular transfer of backward masking under some conditions (Kinsbourne and Warrington, 1962a; Schiller, 1965) suggests the involvement of V1, where signals from both eyes are first processed. Depending on the exact elements constituting the stimulus and the mask, the use of bulk-tissue methods (such as fMRI and EEG) have implicated a variety of brain regions in masking, including lower and higher visual cortical areas (Green et al., 2005; Haynes et al., 2005; Tse et al., 2005), parietal cortex, anterior cingulate, and thalamus (Green et al., 2005). Single unit recordings in macaque monkeys and cats show correlates of masking in retinal ganglion cells (Coenen and Eijkman, 1972), lateral geniculate nucleus (LGN; Coenen and Eijkman, 1972; Macknik and Martinez-Conde, 2004b; Schiller, 1968), V1 (Bridgeman, 1980; Macknik and Livingstone, 1998), V2/V3 (Maeda et al., 2010), and inferotemporal areas (Kovács et al., 1995; Rolls et al., 1999; Rolls and Tovee, 1994). However, none of these studies used perturbations to infer causality. Thus, the circuits determining masking remain largely unknown.

Mice are an essential species for dissecting brain circuits. However, whereas mice have been successfully trained in visual contrast detection (Burgess et al., 2017; Busse et al., 2011), change detection (Garrett et al., 2020; Glickfeld et al., 2013), and orientation discrimination tasks (Andermann et al., 2010; Resulaj et al., 2018), no study has yet demonstrated backward visual masking. A rat study was inconclusive regarding the feasibility of a masking paradigm (Dell et al., 2018; but see Watanabe et al., 2014), particularly because the stimulus regime used in masking (very brief, low contrast stimuli) promoted guessing behavior. We here demonstrate a robust masking paradigm in both mice and humans, with the former permitting the use of genetic tools to dissect the underlying circuit.

## Main text

We modified a 2-alternative choice task (Burgess et al., 2017) in which mice turn a wheel left or right to indicate whether a vertical grating, the target, was presented on the right or left side of a monitor in front of the mouse (Fig. 1A). Performance was quantified by the probability of a response in either direction (*response rate*) and the fraction of responses in the correct direction (*accuracy*). We tested the sensitivity of mice to target duration and contrast (Fig. S1). For all subsequent masking and optogenetic experiments, we chose a target grating of 40% contrast and 17 ms duration. For these parameters, performance was near maximum but approached the steep portion of the psychophysical curve where response rate and accuracy decreased, and reaction times increased.

**Figure 1.**
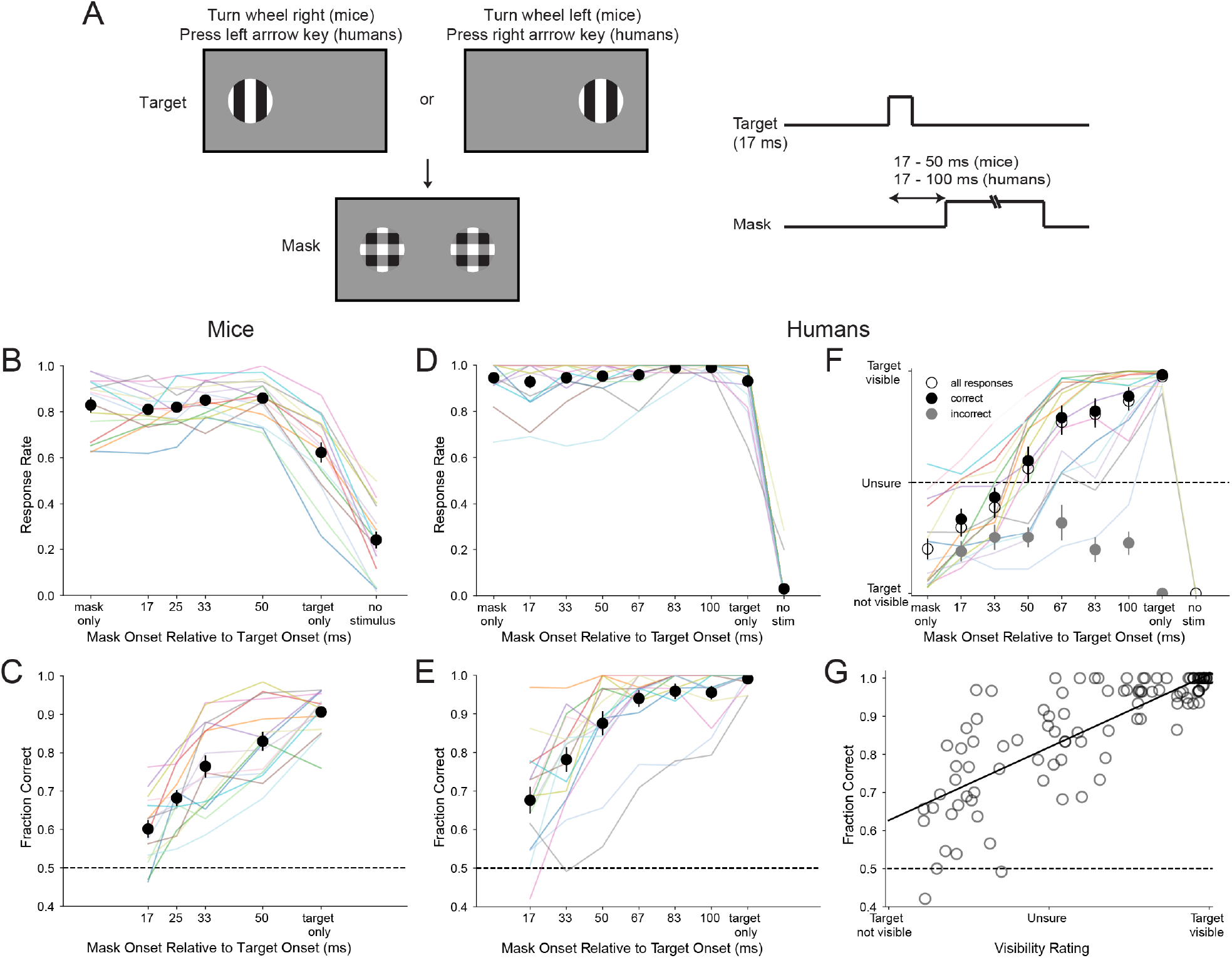
Backward visual masking impairs discrimination of target location. **(A)** A target appeared on the left or right side of a screen in front of mouse or human subjects for 17 ms. Mice earned water rewards by rotating a wheel right or left, respectively. Humans pressed the left or right arrow keys to indicate their choice. On some trials, the target was followed by a mask presented on both sides of the screen with variable onset relative to target onset. Both target and mask were 40% contrast. **(B)** Fraction of trials on which mice responded in either direction (response rate). Colored lines are data from single sessions from individual mice (n=8 wild-type and 8 VGAT-ChR2 mice). Black circles are means across mice; error bars represent standard error of the mean. **(C)** Fraction of responses in the correct direction. Chance accuracy is 0.5 (dashed line). Statistical comparisons between conditions in B-F are shown in Fig. S2. **(D,E)** Same as B and C for 16 human subjects. **(F)** Humans’ report of the subjective visibility of the target side (not visible=−1, unsure=0, visible=1) for all responses (open circles), correct responses (filled black circles, and incorrect responses (filled gray circles). **(G)** Relationship between subjective target visibility and accuracy of human subjects. Each circle represents data for one condition (mask onset or target only) from one subject. The solid, black line is a linear fit of the data (Pearson r=0.81, p<10^−26^).

The mask, a vertical-horizontal plaid of the same size, location, and contrast as the target, was presented for 200 ms on both sides of the screen at variable intervals following target onset (Fig. 1A). Mice had high response rates on masked trials (Fig. 1B). We also evaluated the response rate on trials in which only the mask was presented and on “catch” trials with no visual stimulus; mice never received rewards during these two trial types.

Mice show a monotonic relationship between accuracy and the time of mask onset relative to target onset (Fig. 1C). For the earliest mask onset (17 ms), the mask appeared immediately after the target and accuracy was strongly reduced (no greater than chance for 7/16 mice, binomial test p>0.05). Accuracy recovered to near non-masked levels when mask onset was delayed by 50 ms following target onset. Across mice, there was a significant difference in accuracy across all mask onsets (Fig. S2B). 15/16 mice showed a significant Spearman correlation (r>0.9, p<0.05) between mask onset and accuracy. The response rate on target-only trials was reduced relative to trials with a mask (Fig. 1B, S2A) and to sessions that did not include masks (Fig. S1A,D); yet, high accuracy on target-only trials persisted (Fig. 1C).

These results indicate that the mask disrupts target localization in mice when presented within ~50 ms after target onset. This time course is remarkably similar to backward visual masking in humans and monkeys (see Discussion). However, human masking experiments typically involve discrimination of target features at one location. To determine whether our target location discrimination task produced similar masking in humans, we tested volunteers on the same two-spatial alternative choice task with minor modifications (see Methods). Humans, like mice, were impaired in localizing the target when the mask was presented within 50 ms of the target (Fig. 1D,E, S2C,D). Furthermore, we asked subjects to indicate whether they clearly saw the target (yes, unsure, no). This subjective rating of target visibility decreased with the same time course, revealing a close relationship between performance and perception (Fig. 1F,G, S2E). On error trials, target visibility was judged no different than when the mask was presented alone, regardless of the delay between target and mask onset. Thus, our task produces strong perceptual masking in humans with a similar time course as masking in mice.

Previous experiments in mice using a similar task, but without masking, showed that target detection depended on visual cortex (Burgess et al., 2017). To test whether disruption of target-driven activity in visual cortex is sufficient to explain the effects of masking on behavior, we replaced the mask with bilateral optogenetic inhibition of visual cortex at various times relative to target onset (Fig. 2A). We photo-suppressed excitatory neurons using VGAT-ChR2 mice in which GABAergic interneurons in cortex express channelrhodopsin (Li et al., 2019; Resulaj et al., 2018). Recording of spiking activity across all V1 layers with Neuropixels probes verified that inhibition of putative excitatory neurons was rapid (~12 ms after light onset) and sustained (Fig. 2B, S4A), as previously reported (Bennett et al., 2019; Li et al., 2019; Resulaj et al., 2018). The response rate to the target was reduced to chance levels (Fig. 2C) when visual cortex was inhibited within the time it takes V1 to respond to the target (~45 ms; Fig. S3C,D). The response rate increased when cortical inhibition began as early as 17 ms after the visual response latency (*i.e*., 50 ms after target onset) and rose to near levels observed in the absence of cortical inhibition within ~50 ms of the visual response latency (Fig. 2C, S5A,B). Accuracy on responsive trials was greater than 75% even when the response rate was only slightly above chance (Fig. 2D,E). Unilateral inhibition revealed that the effect of cortical inhibition was specific to the hemisphere contralateral to the target (Fig. S5C). No effect on response rate or accuracy was observed when blue light was delivered above visual cortex in wild-type mice (Fig. S5D,E).

**Figure 2.**
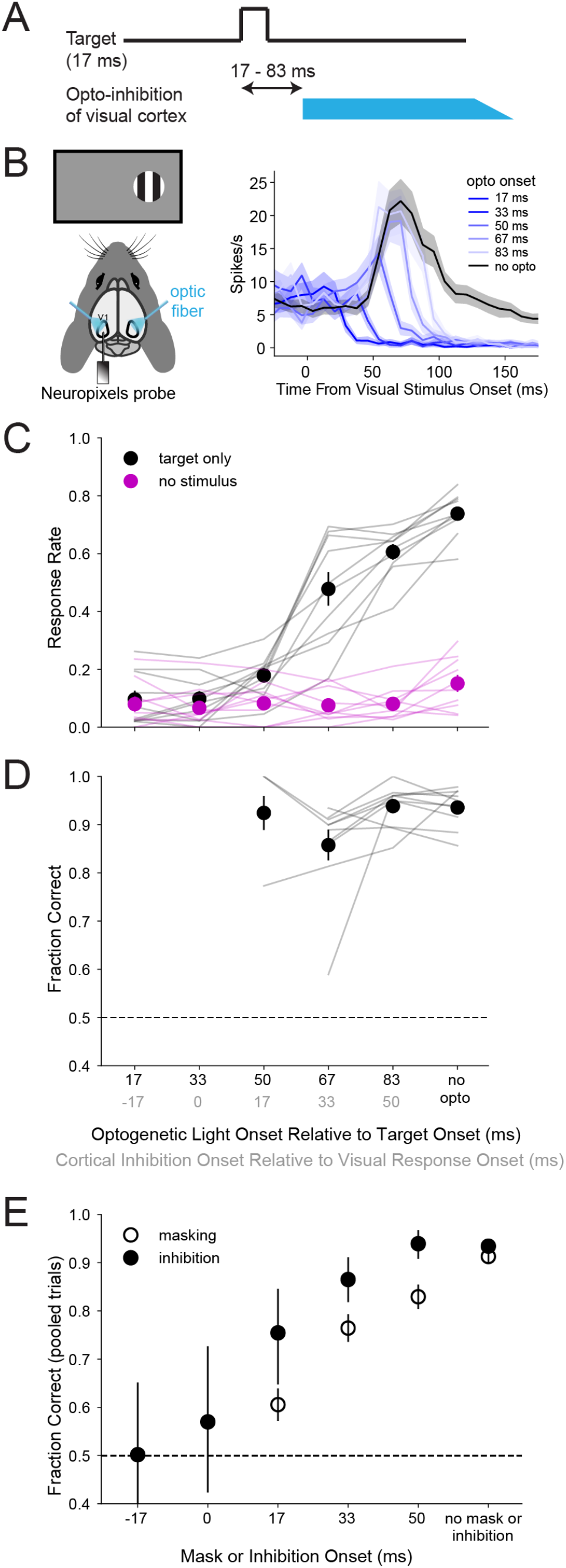
Visual cortical activity during a 50 ms window is required for target detection. **(A)** Bilateral optogenetic inhibition of visual cortex in VGAT-ChR2 mice was varied in time relative to the onset of the target. The target was presented for 17 ms at 40% contrast. The optogenetic stimulus (blue light over V1) persisted for 650 ms after target onset and then linearly ramped off over 100 ms. (B) Mean response of 74 visually-responsive V1 neurons (see Methods) to a contralateral target and the optogenetic stimulus. Neurons that were excited or transiently excited by the optogenetic light were excluded (Fig. S4A). Electrophysiology data was recorded for the experiment shown in Fig. S4B. The behavioral data in C-F were collected during other sessions without electrophysiology. **(C)** Response rate and **(D)** accuracy of mice for different onset times of cortical inhibition. The onset of cortical inhibition relative to the start of the visual response (gray values on x-axis) was estimated by subtracting the visual response latency of ~45 ms (Fig. S3C,D) from the optogenetic light onset and adding the cortical inhibition latency of ~12 ms (Fig. S4A), resulting in an offset of 33 ms between the two sets of x-axis values. Thin lines represent data from single sessions from 10 VGAT-ChR2 mice. Black circles are means across mice; error bars represent standard error of the mean. Statistical comparisons between conditions are shown in Fig. S5A,B. Magenta lines denote “catch” trials with no visual stimulus, used to estimate the chance response rate. Fraction correct values for each mouse in D are only shown for conditions on the x-axis with response rates above chance (>95th percentile of the binomial distribution given the response probability on catch trials and the number of trials; n = 3, 10, 10, and 10 mice for the conditions from left to right). **(E)** Trials were pooled across mice to estimate accuracy when response rates were low (filled circles). Error bars are the 95% confidence interval from the binomial distribution given the fraction correct values and the total number of responsive trials (n = 40, 40, 77, 211, 255, and 1247 trials for the conditions from left to right). For comparison, accuracy on masking trials (Fig. 1) was similarly computed (pooling trials across mice) and plotted (open circles) for corresponding times of mask onset (relative to target onset) or cortical inhibition (relative to cortical visual response latency).

Taken together, our cortical inhibition and masking experiments suggest that a window of target-evoked activity in contralateral visual cortex, lasting approximately 50 ms, is critical for detecting the presence and location of the target. However, masking and optogenetic suppression have distinct effects on behavior. Optogenetically inhibiting bilateral visual cortical activity reduces the probability of responding to the target, with only a modest effect on response accuracy. In contrast, stimulating activity with the bilateral mask (non-informative with respect to target location) decreases the accuracy, but not the rate, of responses. Thus, visual masking is not recapitulated by direct suppression of target-driven spiking in visual cortex.

These observations suggested a model in which the total activity driven in either cortical hemisphere by the target and mask determines the response rate of mice, while the difference in activity between the contralateral and ipsilateral visual cortex determines response accuracy. To explore this idea, we recorded spiking responses of V1 neurons in wild-type mice to the target and mask for the mask onset times shown in Fig. 1B,C. As expected, we observed responses to a contralateral but not ipsilateral target. The response to a contralateral target in the presence of a mask was not a simple sum of the responses to each of these stimuli presented alone (Fig. 3A). Instead, the earliest mask onsets accelerated the spiking response to the mask, while later mask onsets led to a reduced mask response. In correspondence with the response rate of mice (Fig. 1B), the cumulative number of spikes evoked by the target plus mask was larger than the response to the target alone and reached a similar level for all mask onsets of about one spike per neuron on average (Fig. S6A). To estimate the difference in activity across hemispheres (from unilateral recording data), we calculated the difference in cumulative spike counts between trials with a contralateral or ipsilateral target (Fig. 3B,C, S6B). This difference was initially positive, corresponding to spikes evoked by the target on the contralateral side, but quickly converged to zero following onset of the bilateral mask. Thus, on masking trials, spikes informative of target location were only present for very short times, presumably shorter than those used by the mice to perform the task (see below). Consistent with the relevance of interhemispheric comparison, and the requirement of visual cortex for task performance (Fig. 2), the difference in activity across sides was larger when calculated from trials in which the mouse made a correct response compared to error trials (Fig. 3B,C).

**Figure 3.**
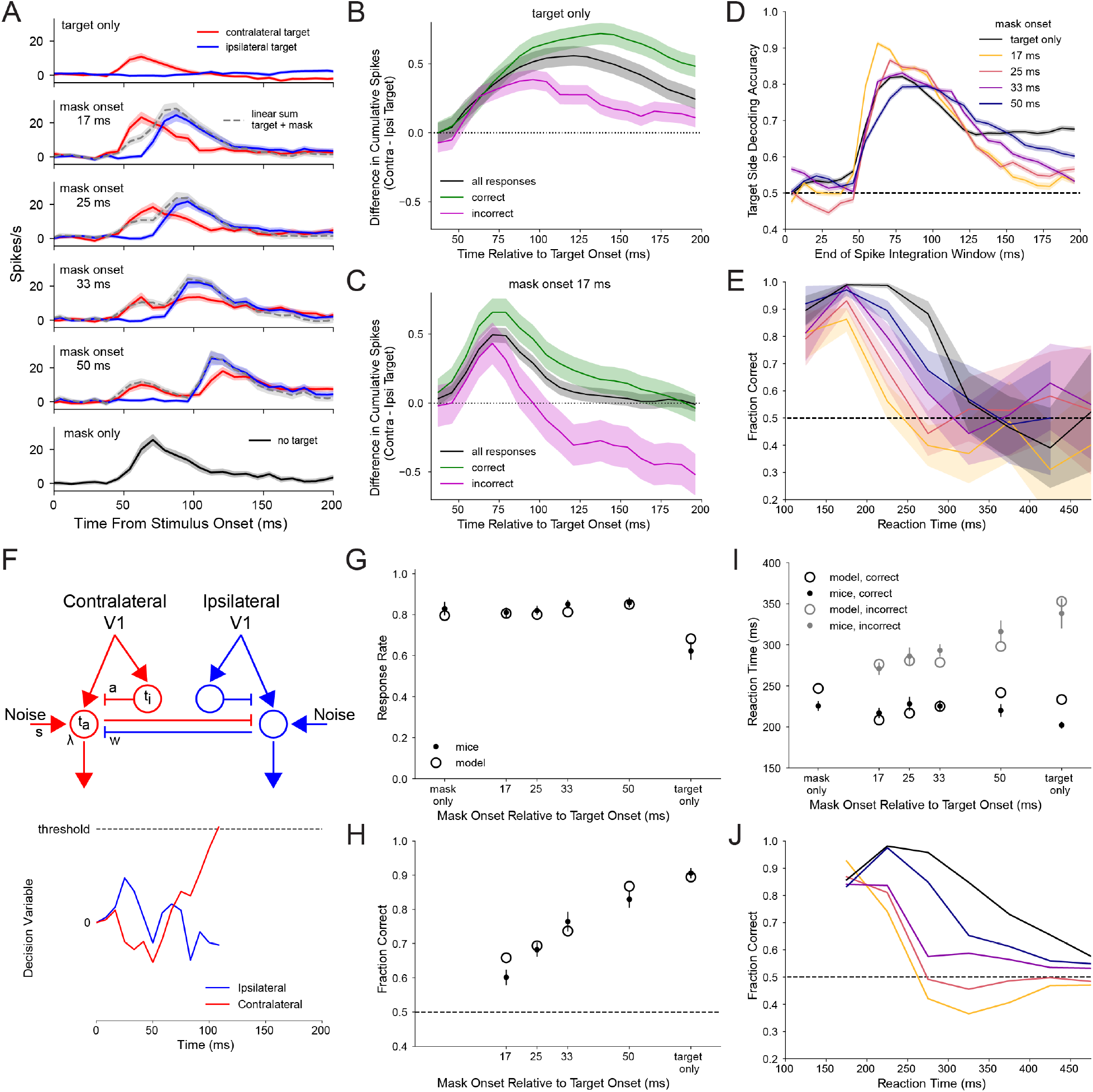
A dual accumulator model using V1 activity is consistent with the performance and reaction times of mice. **(A)** Mean response of V1 neurons (n=58 visually responsive, regularly spiking (RS) neurons from 5 wild-type mice; see Methods) to contralateral (red) and ipsilateral (blue) targets and the mask for the mask onset times in Fig. 1. Mask onset was 0 ms on the x-axis on mask-only trials. Shaded region surrounding each line represents the standard error of the mean. Gray, dashed lines are the sum of responses on contralateral target-only trials and masking trials with an ipsilateral target. An ipsilateral target did not evoke a spiking response on its own (top row). **(B,C)** Difference in cumulative stimulus-evoked spikes per neuron following onset of a contralateral or ipsilateral target for trials with target only (B) or target plus mask onset at 17 ms (C). Values above the dashed, black line at zero are hypothesized to promote correct responses. Spike number was calculated by integrating the trial-averaged post-stimulus time histogram after baseline subtraction for each neuron (as in Fig. S6A). The difference in this value for trials with a contralateral or ipsilateral target was calculated before averaging across neurons. This quantity was calculated for all responses (black line), correct responses (green), or incorrect responses (magenta). The shaded region surrounding each line represents the standard error of the mean. Visually-responsive RS units from wild-type and VGAT-ChR2 mice (Fig. 4, S4, no-opto trials) were combined (n=128) for this analysis. **(D)** Decoding of target location using V1 activity. Linear support vector machines were trained to classify target side using single trial spike counts from the population of V1 neurons. The duration of the integration window, starting at target onset, over which spike counts were calculated varied as indicated by the x-axis. Each line is the mean and standard error across decoders trained and tested using different random subsets of trials (see Methods). **(E)** Fraction correct versus reaction time after pooling reaction times across all mice in 50 ms bins. Shaded region surrounding each line is the 95% confidence interval given the fraction correct values and the number of pooled trials for each bin (the median number of trials across mask onset conditions was 72, 279, 277, 154, 77, 41, 31, and 20 for the time bins from left to right). **(F)** Schematic representation of a dual accumulator model fit to mouse behavior data. Two leaky, mutually-inhibiting accumulators integrate divisively-normalized V1 population activity from contralateral and ipsilateral hemispheres (together with independent noise) in a race to threshold to determine the response direction. The eight free parameters (same values for both “hemispheres”) were the time constants of the accumulator (τ_a_) and the unit mediating divisive normalization (τ_i_), the half-saturation constant for divisive normalization (α), the leakiness of the accumulator (λ), the strength of mutual inhibition (ω), the noise standard deviation (σ), the decision threshold, and the non-decision time (see Methods, Table S2). **(G-I)** Response rate, accuracy, and reaction times of the model (open circles) fit to mouse behavior data (filled circles with error bars representing the standard error of the mean). **(J)** Accuracy versus reaction time for the model, corresponding to E for mice. Panels E and J follow the same colors as D.

To better understand the extent to which V1 population activity is constraining accuracy, we considered an ideal observer with access to V1 activity. We trained linear decoders to classify target location using single trial spike counts of V1 neurons recorded contralateral or ipsilateral to the target. We tested how decoding accuracy depended on the spike integration window by gradually lengthening the time from target onset over which spikes were counted. For short integration windows, the decoder discriminated target location with high accuracy, regardless of whether and when a mask appeared (Fig. 3D). As the integration window extended over longer durations, decoder accuracy declined to levels consistent with the relative accuracy of mice for different mask onsets (Fig. 1C, 3D). Thus, while the initial wave of stimulus-evoked V1 activity is highly informative of target location, spikes evoked by the mask obscure target location over longer integration windows.

Why do mice fail to discriminate the location of the target when followed by a mask if, in principle, they could perform equally well on target-only and masking trials using the earliest wave of V1 activity? Our analysis of V1 population activity suggests that masking occurs when activity is integrated over longer time scales associated with a degraded representation of target location. This predicts that rapid decisions (and corresponding behavioral responses) will tend to be accurate, whereas delayed decisions will more likely be errors. Consistent with this prediction, median reaction times were longer on incorrect trials compared to correct trials for all mask onset times and target-only trials (Fig. S7A-C); the speed of wheel movements, once initiated, did not differ (Fig. S7D,E). Moreover, pooling data across mice revealed an inverse relationship between reaction time and accuracy (Fig. 3E). For each mask onset time, accuracy was highest for the earliest responses and declined to chance with time, but the decline in accuracy began sooner for earlier mask onsets.

Human reaction times differed from those of mice in two ways: (1) they were substantially longer (207 ms on average for mice on target-only trials, versus 595 ms for humans), and (2) there was a strong relationship between reaction times and the time of mask onset (Fig. S7F-H). On correct trials, reaction times were faster with later mask onsets. On error trials, reaction times were similar to when the mask was presented alone, regardless of the delay between target and mask onset. Despite these differences between mice and humans, the accuracy of humans was inversely related to reaction time for each mask onset time (Fig. S7I), as in mice. Hence, both mice and humans “escape” from masking on trials with quick decisions.

Drift diffusion and accumulator models of perceptual decision-making account for performance and reaction times in a variety of tasks (Gold and Shadlen, 2007; Smith and Ratcliff, 2004). We studied whether such models could explain the response rate, accuracy, and reaction times in our visual masking paradigm. To capture the left/right decision, we used a dual accumulator model in which evidence supporting each choice accumulates separately amidst noise towards a threshold; choice is determined by the first accumulator to reach threshold (Fig. 3F). Mouse V1 population activity during trials with a contralateral or ipsilateral target (Fig. 3A) served as the input signal to the two accumulators after divisive normalization (see Methods). We fit the model using a grid search to minimize the sum-of-square error between the response rate, accuracy, and reaction times of the model and mice or humans. The model was able to closely recapitulate the behavior of both mice and humans despite their vastly different reaction times (Fig. 3G-I, S7J-L). This was largely the result of three times greater noise in the accumulators relative to the decision threshold in the mouse model (Table S2), leading to quicker decisions. The relationship between reaction times and accuracy of the models showed the same inverse relationship observed in both mice and humans (Fig. 3J, S7M).

Our data and model suggest that high accuracy on target-only trials can be explained by the relative difference in cortical spiking across the two hemispheres, and that the bilateral mask reduces accuracy by narrowing this difference. The role of visual cortex in backward masking is unresolved (see Discussion). If the critical site where mask-evoked activity disrupts the target signal is in or downstream of visual cortex, as opposed to upstream circuitry (retina, thalamus) or alternative visual pathways (e.g., superior colliculus), our results predict that selectively suppressing mask-evoked activity in visual cortex should rescue response accuracy (*i.e*., prevent backward masking). To test this prediction, we inhibited cortical activity bilaterally at various times relative to the mask (using the earliest and most effective mask onset time, 17 ms; Fig. S4B). For the earliest onset of visual cortex inhibition that allowed a response rate greater than chance, response accuracy on masking trials was increased to a level not significantly different from target-only trials (Fig. 4A,B, S8A,B). This improvement in accuracy decreased monotonically as cortical inhibition was delayed relative to target onset, allowing more mask-evoked spikes. The optogenetic light stimulus had no effect in wild-type mice (Fig. S8C,D). Thus, appropriately timed inhibition of visual cortex that preserves the earliest target-evoked spikes but suppresses mask-evoked activity prevents backward visual masking, revealing an important role for visual cortex in this process.

**Figure 4.**
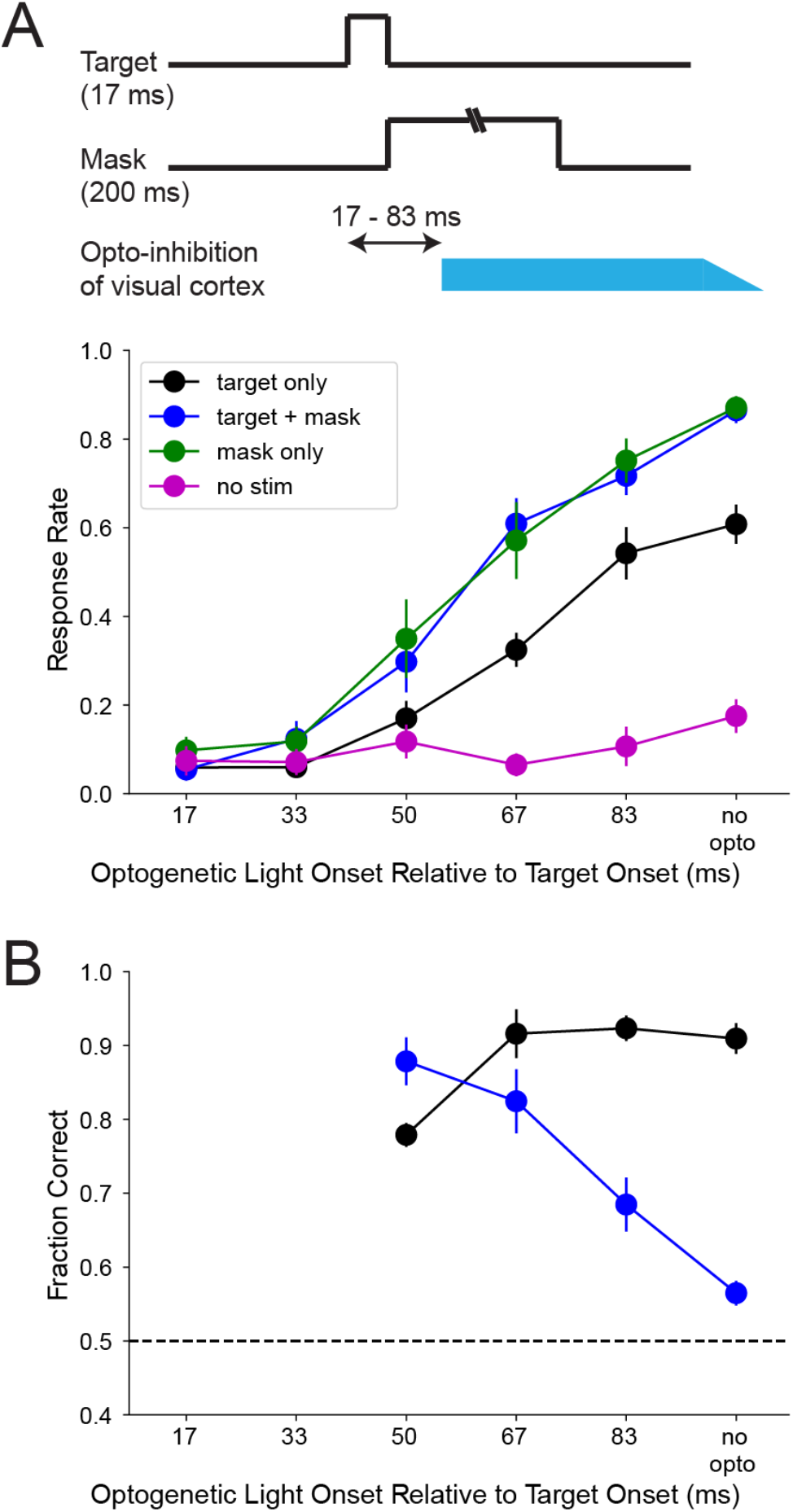
Optogenetic inhibition of mask-evoked activity in visual cortex rescues accuracy of behavioral responses. **(A)** Response rate and **(B)** accuracy for sessions during which the onset of bilateral optogenetic silencing of visual cortex was varied in time relative to the onset of the target and mask. The target stimulus was presented for 17 ms and, on masking trials, was immediately followed by the mask. Circles are means from 8 VGAT-ChR2 mice; error bars represent standard error of the mean. Statistical comparisons between conditions are shown in Fig. S8A,B.

## Discussion

We successfully developed a backward visual masking paradigm to explore the precise timing and circuits underlying visual perception in a model organism amenable to mechanistic circuit dissection. Our masking paradigm uses a variant of a 2-alternative choice task developed by the International Brain Laboratory to study spatial decision-making (Burgess et al., 2017; IBL et al., 2021). We modified the task to use targets of very short duration presented at one of two possible locations, combined with a bilateral mask. Variations of this task could study other forms of masking, such as metacontrast masking (Alpern, 1953) in which the spatial location of the mask does not overlap the target.

The target was susceptible to masking for ~50 ms after stimulus onset in both mice and humans, similar to the timescale of masking previously seen in humans (Alpern, 1953; Bachmann and Francis, 2013; Bacon-Macé et al., 2005; Breitmeyer and Ogmen, 2006; Dehaene, 2014; Eriksen and Lappin, 1964; Kinsbourne and Warrington, 1962b; Kolers, 1962; Schiller, 1965; Sperling, 1965) and monkeys (Kovács et al., 1995; Macknik and Livingstone, 1998; Rolls et al., 1999). The effect of the mask decreased monotonically with duration from target onset, typical for our parameters in which the mask spatially overlaps the target and is presented for a relatively longer duration (Bachmann and Francis, 2013; Breitmeyer and Ogmen, 2006). This shared phenomenology across mammalian species suggests our masking paradigm can be a powerful tool for dissecting circuit mechanisms.

By silencing visual cortical activity at precise time points following the mask, we could ameliorate the behavioral effects of masking in mice. This demonstrates a crucial role of visual cortex and indicates that target-mask interactions in the retina, thalamus, and/or superior colliculus are insufficient on their own to generate masking in our task. In human studies, transcranial magnetic stimulation (TMS) of visual cortex following the mask can increase target detection (Amassian et al., 1993; Ro et al., 2003). However, subjects also reported seeing the target on trials when the target was not presented, perhaps due to phosphene generation (Ro et al., 2003) or incomplete suppression of the mask (which was then perceived as the target). Hence, the exact cortical perturbation caused by TMS in masking experiments is not well understood (Bachmann and Francis, 2013). In contrast, we directly measured and suppressed mask-evoked spiking in visual cortex in a well-controlled manner to show it is necessary for backward masking.

Our data demonstrate distinct effects of bilateral cortical silencing (via optogenetic inhibition) and bilateral masking – the former primarily reduces the probability of responding while the latter affects primarily the accuracy of responses.

Together, these data are compatible with the hypothesis that the initial target-evoked spike in visual cortex can be sufficient for an ideal observer to decode target location with high accuracy, consistent with results in humans (Thorpe et al., 1996), monkeys (Kovács et al., 1995; Rolls and Tovee, 1994) and mice (Resulaj et al., 2018). Indeed, our decoding analysis indicates that, on average, neurons respond to the target stimulus with around one action potential in the relevant time window. In the presence of a mask, the target location becomes ambiguous as downstream decision stages integrate over longer durations. This likely reflects sensory integration mechanisms and decision thresholds tuned for noise reduction rather than resistance to masking. The brain regions performing this temporal integration of visual cortical signals remain unknown.

Interestingly, even in the face of a mask, accuracy on the fastest reaction time trials can be very high (~90%). Previous human studies likewise show that the fastest reactions can evade masking in tasks using either saccadic (Crouzet et al., 2014) or manual responses (Bacon-Macé et al., 2005; Van Rullen and Koch, 2003). However, absolute accuracy in these studies is much lower on the fastest reaction time trials (e.g. ~60% in Crouzet et al. 2014), which could reflect shorter neural processing times for the simpler spatial discrimination in our task. Escape from masking has been interpreted as evidence that the mask disrupts reentrant feedback processing while sparing the initial feedforward wave (Crouzet et al., 2017, 2014; Van Rullen and Koch, 2003). Our dual accumulator model suggests that the lack of integration of feedforward target and mask signals before reaching a decision threshold can also explain the similar accuracy on fast reaction time trials for target-only and masking trials. Similar models have been successful in a variety of contexts (Gold and Shadlen, 2007; Smith and Ratcliff, 2004), but rarely have been applied to visual masking (Ratcliff and Rouder, 2000; Vorberg et al., 2003), and never with a neuronally constrained model. Our work suggests that such models can help bridge perceptual decision-making and backward visual masking. Although our task involved discriminating activity between spatially segregated populations of neurons representing different target locations, the dual accumulator model can similarly apply to spatially co-mingled neurons differentially tuned to features of a target, appearing at a fixed location, such as orientation or direction of motion.

In humans, visual masking goes hand in hand with a loss of perceptual visibility of the target. Indeed, in our two-location masking paradigm we find that human subjective report of visibility corresponds closely with behavioral performance. Whether the same phenomenology exists in mice remains unanswered for now. Future studies can adapt our task to include measures of subjective confidence that could begin to address this question in mice (Kepecs et al., 2008; Kepecs and Mainen, 2012).

## Acknowledgements

We thank the Allen Institute founder, Paul G. Allen, for his vision, encouragement, and support. We thank the Tiny Blue Dot Foundation for providing funding for this study. We thank the Neurosurgery and Behavior team for help with mouse surgeries. We thank Sarah Naylor for support with program management. We thank Shannon Reynolds for help with administration of the animal and human protocols. We thank Masataka Watanabe and Doug Ollerenshaw for discussion of masking paradigms in rodents.

## Author contributions

SDG, CB, CK, and SRO conceived the project. SDG, CS, CB, and SRO developed the mouse and human experimental methodology. SDG and CS performed the experiments. SDG and CB analyzed the data. SM provided advice on modeling and data analysis. CK and SRO supervised all aspects of the project. All authors contributed to writing the manuscript.

## Competing interests

The authors declare no competing interests.

## Methods

### Mice

All experiments were performed at the Allen Institute using protocols approved by the Allen Institute’s Institutional Animal Care and Use Committee. We used 8 wild-type mice (C57BL6J; 3 female, 5 male) and 10 VGAT-ChR2 mice (6 female, 4 male) (Zhao et al., 2011) for experiments (Table S1). Two of the VGAT-ChR2 mice were used for the cortical inhibition experiment shown in Fig. 2 but not for masking experiments. An additional 5 wildtype mice and 3 VGAT-ChR2 mice were not used for experiments due to poor performance during initial training on the target location task without masking.

Mice were single-housed and maintained on a reverse 12-hour light cycle. All experiments were performed during the dark phase. Prior to the start of training on the behavioral task, mice were anesthetized with isoflurane and a headpost was attached to the skull with clear dental acrylic. After one week of recovery, mice were provided daily with an amount of water required to maintain 85% of their initial body weight, with continued unrestricted access to food. Mice were 67-145 days old (median 89) at the start of training.

### Humans

Human participants were Allen Institute employees who were not otherwise involved in the study. Protocols were approved by the Institutional Review Board and participants were required to provide written, informed consent. Volunteers did not receive compensation. Preliminary data used to set parameter values was collected from three volunteers. The final data set was collected from 16 volunteers.

### Mouse Behavior Task

The behavior task was adapted from Burgess et al. 2017. Mice were head-fixed on a platform with their front paws resting on a 6 cm diameter wheel. Visual stimuli were controlled using Psychopy version 3.2.4 (Peirce, 2009) and presented at 120 frames/s on a monitor (ASUS VG248QZ; 1920 x 1080 pixels, 53.3 cm wide) centered 21.6 cm in front of the mouse. The bottom right corner of the screen (50 x 50 pixels) flipped between black and white each frame and was covered with a photodiode. The signal from the photodiode was used to determine frame times. The target stimulus was a circular vertical grating (25° diameter, 0.08 cycles/°) positioned at the center of either the left or right half of the screen (31.7° horizontal and 0° vertical from the mouse). The mask was a pair of vertical-horizontal plaids of the same size, spatial frequency, and position (both left and right sides) of the target. The outer 8% of the target and mask were cosine blurred. The luminance of the blank/gray screen was 60 cd/m^2^. The target was 100% contrast during training and for testing sensitivity to target duration (Fig. S1A-C), variable contrast for testing contrast sensitivity (Fig. S1D-F), and 40% contrast for masking and cortical inhibition experiments. The mask was 40% contrast.

Each trial began with a blank, gray screen for a fixed 3 s followed by a variable duration drawn from an exponential distribution (mean 1 s, max 5 s) before the target appeared. Mice were rewarded with a drop of water (~2.5 μL) delivered from a lick spout below the mouth for rotating the wheel left when the target was on the right side of the screen, and vice versa. During initial training, the target persisted on the screen and movement of the wheel was coupled to movement of the target in the same direction. Wheel movement during the first 133 ms after target onset was ignored. The movement threshold for reward was initially 3-10 mm along the circumference of the wheel (adjusted for each mouse at the trainer’s discretion). The gain between wheel and target movement was set such that the target was at the center of the screen when the reward threshold was reached. Mice were initially given a response window of up to 30 s after target onset to move the wheel past threshold. Movement past threshold in either direction (correct or incorrect) during the response window ended the trial. Lack of movement past threshold within the response window was deemed “no response.” Trials with either an incorrect or no response were repeated up to 5 times during training.

As mice learned to turn the wheel in response to the target, the response window was gradually decreased to a final value of 650 ms from target onset. Three additional task features were introduced: (1) “catch” trials (15% of trials during training) with no visual stimulus or reward were used to monitor the rate of non-visually driven movements past threshold during the response window; (2) a quiescent period in which movement of the wheel >1mm in either direction during the last 500 ms before target onset restarted the variable gray screen period with a new random duration; and (3) incorrect responses were followed by a 1 s white noise burst and an additional 3-6 s timeout period (during training only). For some training sessions, the probability that the target appeared on the right or left was biased from the normally equal probability to correct movement biases for individual mice.

After mice were consistently performing the task at >90% accuracy with wheel and target movement coupled (16-51 training sessions; median 32), mice were trained to respond to flashed target stimuli with fixed position and duration of 33-200 ms for 1-5 sessions. The reward threshold was reduced to 2 mm. Following these sessions, sensitivity to target duration and contrast were tested (Fig. S1). During these experiments, the set of possible target durations (8, 17,33 and 100 ms) or contrasts (20, 40, 60, and 100%), and target sides (left, right), were randomly sampled before repetition. The catch trial probability was set to 11% such that there were approximately equal numbers of target-left and target-right trials (for each duration or contrast) and catch trials (Table S1).

For masking experiments (Fig. 1B,C), on each trial there was a 60% probability that either the target (17 ms, 40% contrast) was followed by the mask (40% contrast, 200 ms) after a variable interval relative to target onset, or the mask was presented alone (mask-only trials). The set of possible mask onset times (17, 25, 33, and 50 ms relative to target onset) and target sides were sampled in random order before repetition. The other ~40% of trials were target-only trials or catch trials with no visual stimulus. Rewards were never delivered on mask-only or catch trials. The probability that a trial included no target presentation (mask-only and catch trails) was set to 11% such that there were approximately equal numbers of target-left and target-right trials (for each mask onset) and no-target trials (Table S1).

### Human Behavior Task

The human version of the behavioral task was identical to the mouse version unless otherwise noted. Participants rested their chin and forehead against a tabletop chin rest and were instructed to fixate on a cross at the center of the screen. Eye position was not monitored. The blank screen was 28 cd/m^2^. The target stimulus (2° diameter, 2 cycles/°) was presented 15.3° to the right or to the left of the fixation cross. Target and mask were 40% contrast. Mask onset times were 17, 33, 50, 67, 83, and 100 ms after target onset. The mask persisted throughout the response window (2.5 s). The mask was presented on ~75% of trials, and the other ~25% of trials were target-only or catch trials. Participants were instructed to press the right or left arrow key to indicate whether they saw the target on the right or left, respectively, within 2.5 seconds of target onset. There was no feedback on the accuracy of responses. At the end of the response window, a question immediately appeared on the screen asking “Did you see the target side? No=1, Unsure=2, Yes=3” and subjects had as much time as they required to answer by pressing the 1, 2, or 3 key. Participants practiced for a few trials to make sure they understood the task. They then completed 260 trials, which took 28-36 minutes (Table S1).

### Optogenetic inhibition of visual cortex

For optogenetic inhibition of visual cortex in VGAT-ChR2 mice, or control experiments in wild-type mice, optical fibers (200 μm diameter tip) were placed bilaterally over the skull 2.7 mm lateral and 0.5 mm anterior of lambda (the skull was covered by a thin layer of clear dental acrylic during surgery). A black plastic cone was positioned above the headpost to block stray light. For all trials, there was 60% probability that blue light (470 nm, 2.5 mW) was delivered from one or both of the optical fibers. The set of possible optogenetic light onset times (17, 33, 50, 67, and 83 ms relative to target onset; Figs. 2 and 4) or brain hemispheres (Fig. S4C) and target sides were sampled in random order before repetition. The blue light persisted until the end of the behavioral response window (650 ms after target onset) and then linearly ramped off over 100 ms. The probability that a trial included no target presentation (mask-only or catch trails) was set to 33% such that there were approximately equal numbers of target-left, target-right, and no-target trials for each optogenetic light onset (Table S1).

### Electrophysiology

Putative V1 recordings were made during separate experiments/days after the behavioral sessions analyzed for Figures 1, 2, and 4. The day before recording, or the same day at least 4 hours before recording, mice were anesthetized with isoflurane and a 1 mm diameter craniotomy was made 2.7 mm lateral and 0.5 mm anterior of lambda above the left hemisphere. During recordings in VGAT-ChR2 mice, the left optical fiber was placed above the skull at the medial edge of the craniotomy. Neurons were recorded with Phase 2 Neuropixels probes (128 channels arranged in two columns, with 20 μm between each recording site) (Jun et al., 2017) throughout all cortical layers. Data were acquired at 30 kHz using the Open Ephys acquisition board and GUI (Siegle et al., 2017).

At the beginning of each recording, receptive fields were coarsely mapped by presenting a 25° diameter vertical-horizontal plaid (100% contrast, 0.08 cycles/°) centered at one of 9 equally spaced positions that formed a grid across the right half of the display. The stimulus was presented for 50 ms at all 9 positions in random order (with 800 ms between presentations) before repetition. After 25-30 repeats, spikes were detected across all channels using an amplitude threshold, and post-stimulus time histograms were calculated for each stimulus position. If the peak response was greatest at the center position (where the target/mask would be presented), we continued to record during the masking task. Otherwise, we moved the probe and repeated this process, or ended the experiment.

### Analysis of behavior data

Data for each experiment (Fig. 1,2,4) are based on a single session from each mouse. Trials were excluded if an unexpectedly long frame interval disrupted the timing of stimulus onset or duration (0.4% of trials across all sessions), the mouse moved the wheel >1 mm during the first 125 ms after target onset (3.4% of trials), or the mouse did not respond during the previous 10 trials (not including catch trials or trials with optogenetic inhibition; 1.2% of trials). Data from target-left and target-right trials were pooled. Reaction times were calculated as the last time point after stimulus onset that wheel displacement was greater than a movement initiation threshold of 0.2 mm before crossing the reward threshold (2 mm) in the same direction. Movement speed was calculated as the difference between the reward and movement initiation thresholds (1.8 mm) divided by the difference between the times these thresholds were crossed.

For human subjects, reaction time was the time that a key press was detected relative to stimulus onset. Answers to the question “Did you see the target side?” were coded as −1 (“no”), 0 (“unsure”), or 1 (“yes”) for the quantification shown in Fig. 1F,G.

### Analysis of electrophysiology data

Spikes were sorted automatically by Kilosort2 (Stringer et al., 2019). Putative single units and multi-units were accepted for further analysis if the average firing rate over the entire session was greater than 0.1 spikes/s and the majority of spikes were from a single neuron (false-positive rate less than 0.5) as estimated by comparing the rate of refractory period (1.5 ms) violations to the overall spike rate (Fig. S3A) (Hill et al., 2011; Siegle et al., 2021). Units were defined as fast-spiking (FS; putative inhibitory interneurons) if the peak-to-trough duration of the average spike waveform was less than 0.4, or otherwise as regular spiking (RS; putative pyramidal neurons; Fig. S3B). Time to first spike (Fig. S3D) (Siegle et al., 2021) for each unit was calculated as the median across trials of the time of the first spike 30 ms or later after visual stimulus onset; trials with no spike within 150 ms of stimulus onset were excluded.

Post-stimulus time histograms (PSTHs) of neural activity aligned to visual or optogenetic stimulus onset were calculated using 8.33 ms bins (the monitor frame duration). Units were considered visually responsive if they had a significant response to either the target or the mask. A response was considered significant if (1) the peak of the PSTH after visual stimulus onset was greater than 5 times the standard deviation of the PSTH during the last 500 ms before stimulus onset, and (2) a Wilcoxon signed-rank test comparing the spike rate 40-100 ms after stimulus onset and the spike rate before stimulus onset resulted in a p-value less than 0.05.

Based on responses to the optogenetic light stimulus (Fig. S4A), units recorded in VGAT-ChR2 mice (n=267 units from 4 mice) were classified as (1) excited if the mean of the PSTH 100-500 ms after light onset was greater than the standard deviation of the PSTH during the last 500 ms before light onset; (2) transiently excited if the peak of the PSTH during the first 100 ms after light onset was greater than 5 times the standard deviation of the PSTH before light onset; or (3) inhibited if there was no transient excitation and the mean of the PSTH 100-500 ms after light onset was less than the standard deviation of the PSTH before light onset. 74 visually-responsive units that were not excited or transiently excited by the optogenetic light stimulus were analyzed for Fig. S4B.

58/276 units from 5 wild-type mice were visually-responsive RS units and contribute to the data shown in Fig. 3A,D and S6. Visually-responsive RS units from wild-type and VGAT-ChR2 mice (excluding trials with optogenetic light delivery) were combined (n=128) for the analysis comparing activity on correct and incorrect trials shown in Fig. 3B,C.

For the decoding analysis shown in Fig. 3D, neurons were pooled across mice. For each mouse/session, trials for each target side (contralateral or ipsilateral to the recording) and mask onset condition were evenly spilt at random into training and testing groups. We then created 100 training and testing pseudo-trials for each condition by randomly selecting a trial from the training or testing group for each neuron. A linear support vector machine (SVM) was trained to classify the target side for each pseudo-trial from the training group using the spike count from each neuron as input. Decoder accuracy was then determined from the decoder’s performance on the pseudo-trials from the testing group. This process was repeated for 100 separate splits of training and testing data to determine the average decoder accuracy for each duration over which spikes counts were quantified (x-axis of Fig. 3D).

### Dual accumulator model

The model of mouse and human behavior shown in Fig. 3F is a race-to-threshold between two accumulators integrating activity from the population of V1 neurons on trials with a contralateral or ipsilateral target, respectively. The output of the accumulators, L and R, is determined by the following equations,

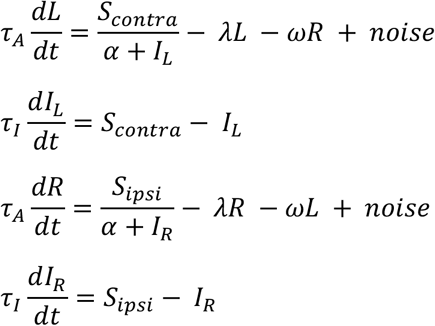

S_contra_ and S_ipsi_ are trial-averaged V1 population activity on trials with a contralateral or ipsilateral target (Fig. 3A) and provide input to the accumulators L and R, respectively. I_L_ and I_R_ integrate the same V1 activity and divisively normalize the input to L and R, respectively. The 8 free parameters are the time constants of L and R (τ_A_) and I_L_ and I_R_ (τ_I_), the half-saturation constant for divisive normalization (α), the leakiness of the accumulators (λ), the strength of mutual inhibition (ω), the noise standard deviation (σ), the decision threshold, and the non-decision time accounting for the delay between decisions and movement. These parameters were the same for both accumulators, but the input and noise on each time step was independent. The time step, dt, was 8.33 ms. On each trial, the first accumulator to reach threshold determined the decision of the model (contralateral or ipsilateral target). If neither accumulator reached threshold the outcome was “no response.” The model parameters were fit via a brute force grid search to minimize the sum-of-square error between the response rate, accuracy, and reaction times of the model and mice or humans. Model error was normalized by the standard error of the data. Values of the fit parameters are given in Table S2.

### Statistics

Statistical tests are described in the main text and/or figure legends.

---

**Figure S1.**
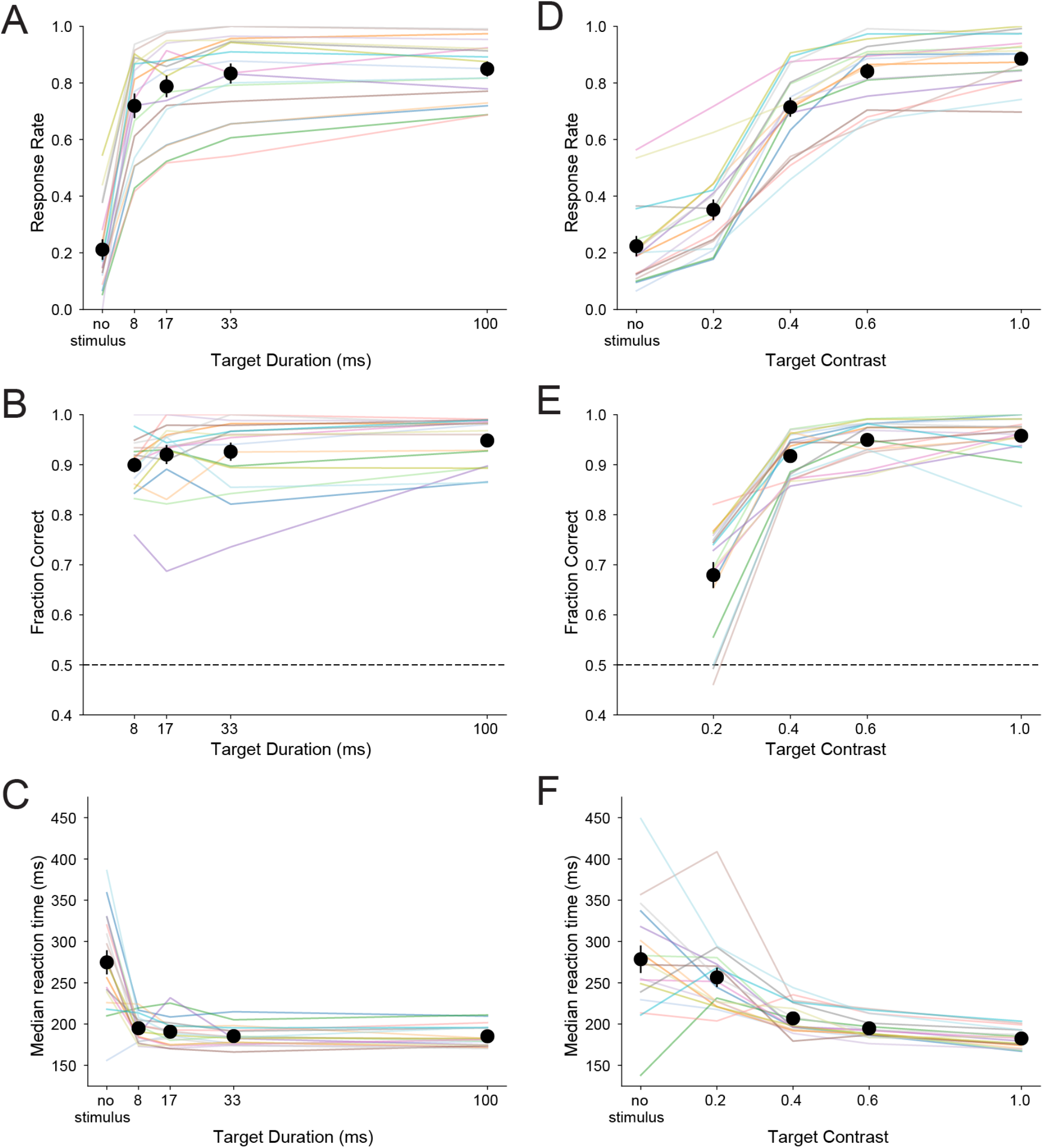
Sensitivity of task performance in mice to target duration **(A-C)** and contrast **(D-F)** in the absence of a mask. Target contrast was 100% for experiments varying target duration (A-C). Target duration was 17 ms for experiments varying target contrast (D-F). Colored lines are data from single sessions from individual mice (n=8 wild-type mice, 10 VGAT-ChR2 mice). Black circles are means across mice; error bars represent standard error of the mean.

**Figure S2.**
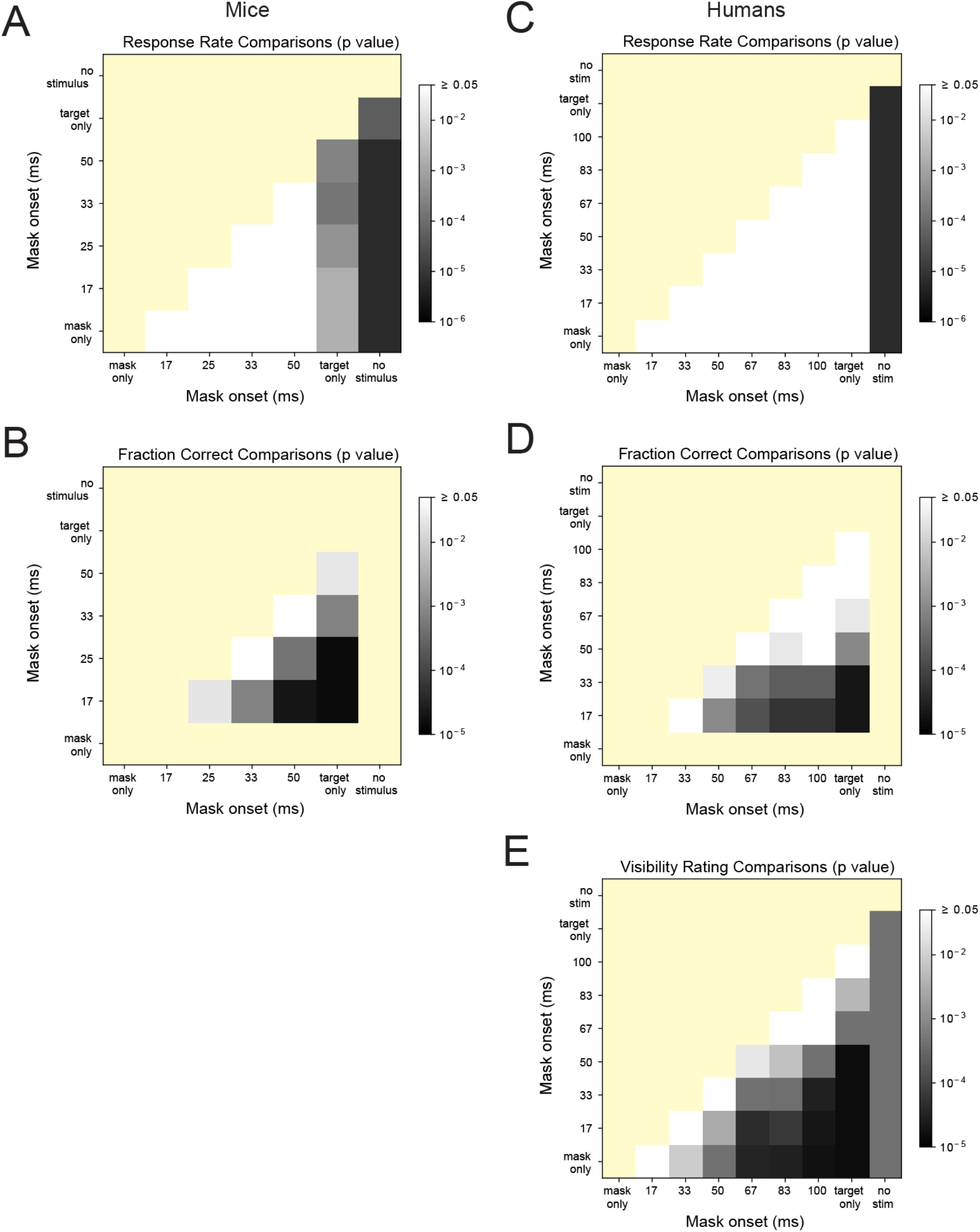
**(A,B)** p-values from comparison of mouse response rates (A) and accuracy (B) for the conditions shown in Fig. 1B,C. Distributions were compared with a Kruskal-Wallis test followed by pairwise Wilcoxon rank sum tests. p-values were corrected for multiple comparisons using the Benjamini-Hochberg method. Greyness scale is log_10_ based. Non-significant p-values (≥0.05) are white. Redundant or non-tested comparisons are yellow. **(C-E)** Same as A and B for human subjects (Fig. 1D-F), additionally showing comparisons of subjective visibility ratings (E).

**Figure S3.**
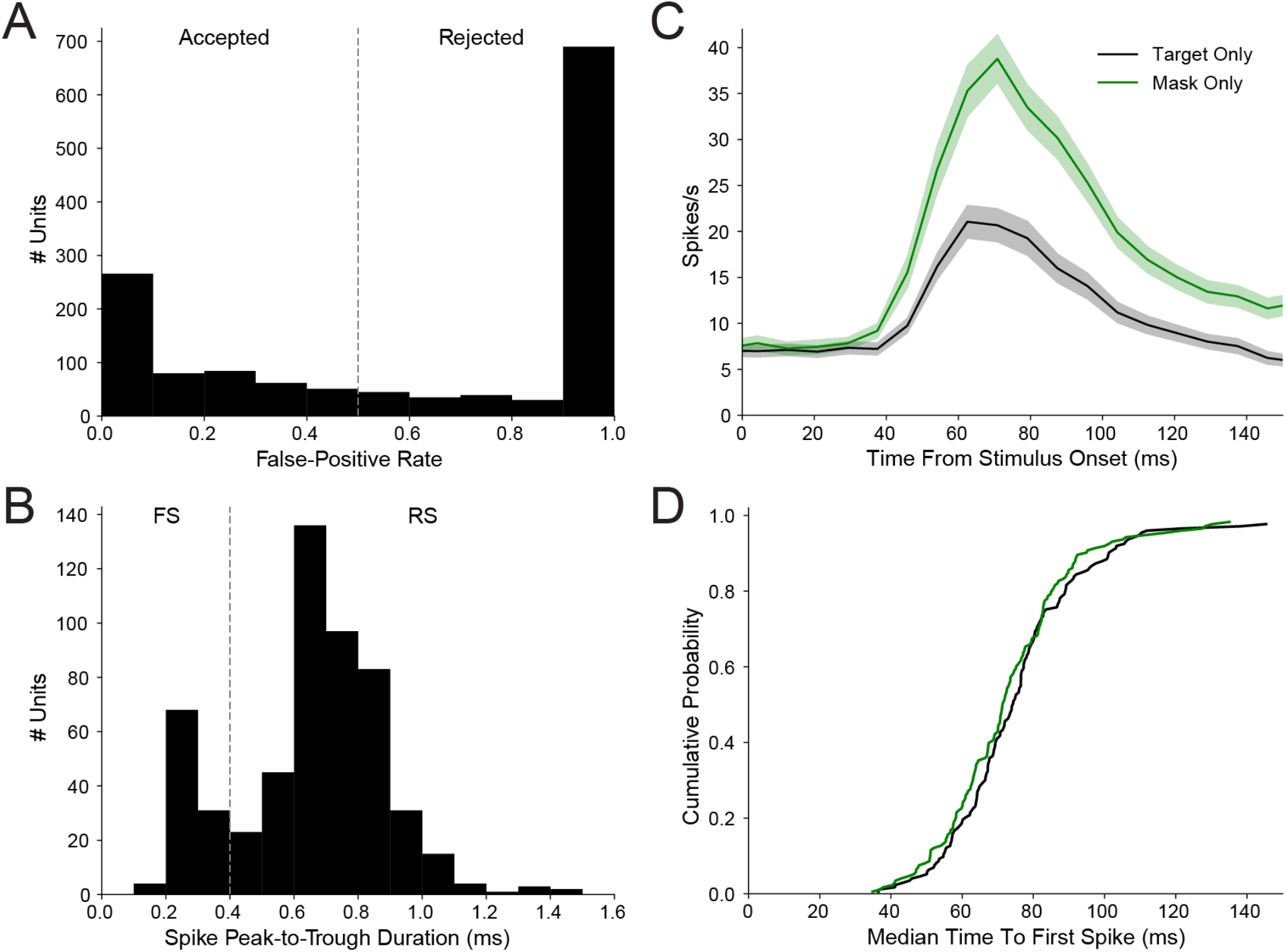
**(A)** False-positive rate for spikes recorded from 1382 units from 5 wild-type and 4 VGAT-ChR2 mice. The false-positive rate was defined as the refractory-period violation rate (number of spikes within 1.5 ms refractory periods divided by the total duration of refractory periods) divided by the total spike rate (see Methods). Units with a false-positive rate greater than one by this definition are displayed at one in the histogram. 543 units with false-positive rates less than 0.5 were considered for further analysis in this study. **(B)** Peak-to-trough duration of the average spike waveforms (n=543). Units with peak-to-trough duration less than 0.4 were classified as fast spiking (FS; putative inhibitory interneurons); all other units were classified as regular spiking (RS; putative pyramidal neurons). **(C)** Average firing rate of 173 visually responsive units (see Methods) following presentation of a contralateral target (17 ms duration) or bilateral mask (200 ms duration) starting at 0 ms on the x-axis. Shaded area is the standard error of the mean. **(D)** Cumulative distribution of the median time to first spike (see Methods) of the 173 visually responsive units. There was no significant difference in the time to first spike between target-only (median across units 73.9 ms) and mask-only (71.3 ms) trials (p=0.22, Wilcoxon rank sum test). There was also no significant difference between the time to first spike of RS (n=128) and FS (n=45) units (target only: 75.1 vs. 68.1 ms, p=0.08; mask only: 71.4 vs. 68.9 ms, p=0.39), which were combined for the data plotted.

**Figure S4.**
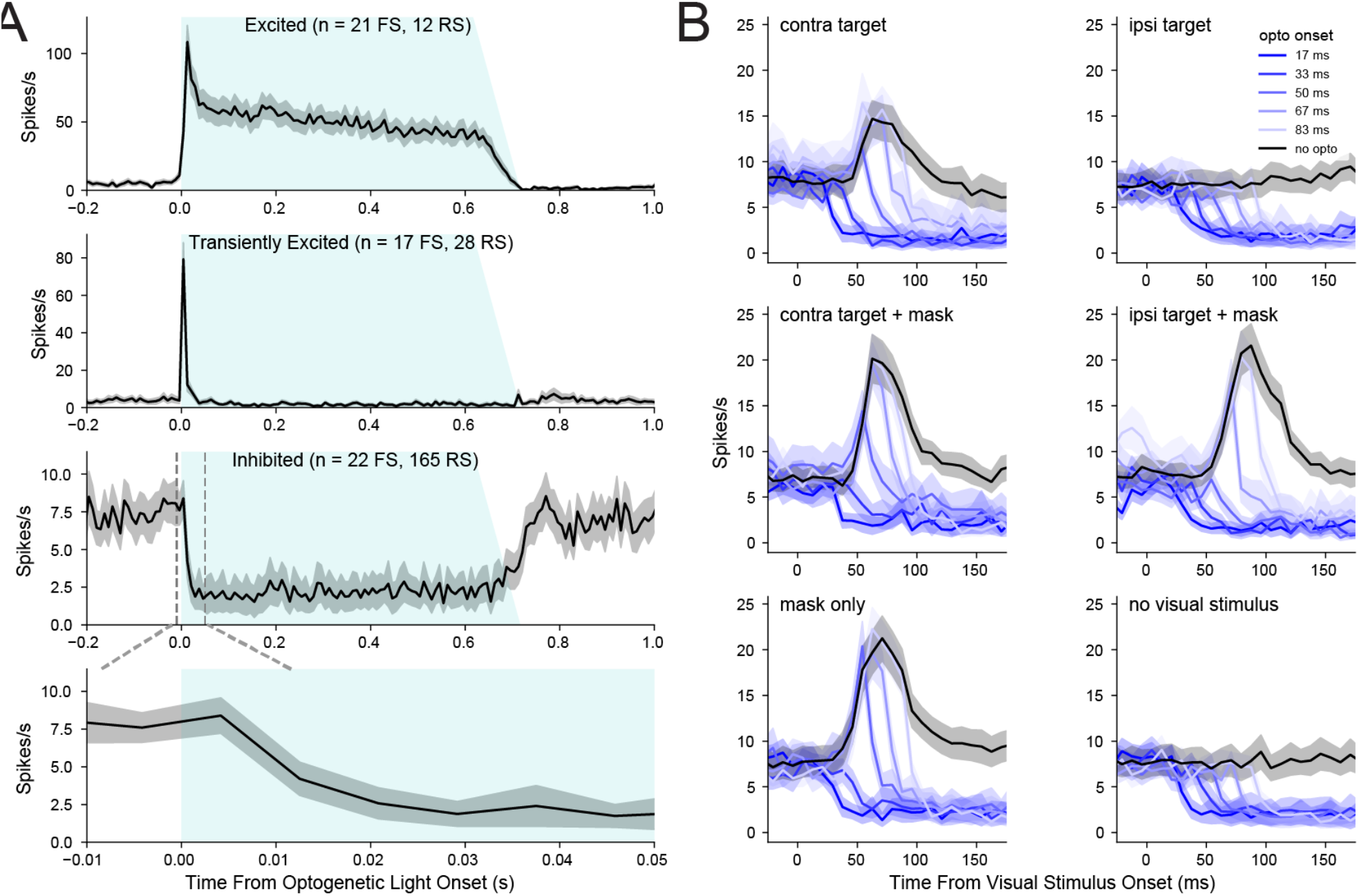
**(A)** Mean response of V1 neurons from 4 VGAT-ChR2 mice to stimulation with blue light. Gray, shaded region indicates standard error of the mean. Fast-spiking (FS, putative interneurons) and regular spiking (RS, putative pyramidal neurons) were defined by the peak-to-trough duration of their spike waveforms. Neurons were classified as excited, transiently excited, or inhibited (see Methods). Two neurons were non-responsive (not shown). The bottom panel expands the time scale (indicated by dashed lines) of the inhibited cells. **(B)** Mean response of 74 visually-responsive V1 neurons (see Methods) from the same mice in A to the visual and optogenetic stimuli used for the behavior data in Fig. 4A,B. Neurons that were excited or transiently excited by the optogenetic light were excluded. Optogenetic light onset was varied relative to the onset of the target presented contralateral or ipsilateral to the recorded neurons. On a subset of trials, the target (17 ms duration) was immediately followed by a bilateral mask for 200 ms.

**Figure S5.**
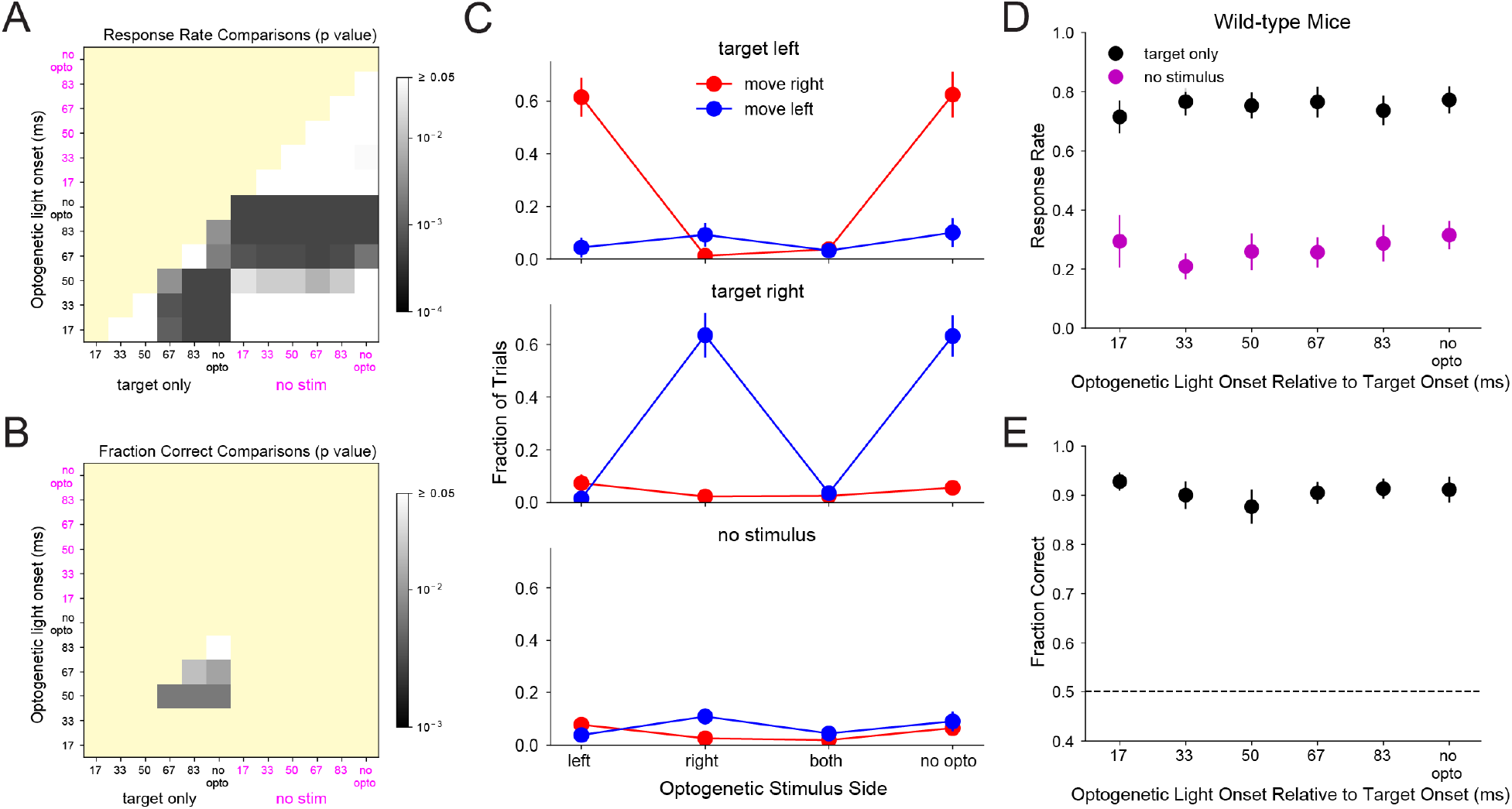
**(A,B)** p-values from comparison of response rates (A) and accuracy (B) for the conditions shown in Fig. 2C,D, using statistical procedures described in the legend for Fig. S2A,B. **(C)** Effect of unilateral cortical inhibition on target detection (n=8 VGAT-ChR2 mice). The inhibited hemisphere(s) are indicated on the x-axis. Optogenetic light onset was 17 ms before target onset. The top plot shows, for trials on which the target was left, the fraction of trials that mice moved right (red, correct) or left (blue, incorrect). The sum of the red and blue data points for each condition is the response rate. The middle and bottom plots show similar data for target-right and no visual stimulus trials, respectively. **(D,E)** Effect of optogenetic light stimulus on response rate and accuracy in wild-type mice (n=8) for the same conditions shown in Fig. 2C,D.

**Figure S6.**
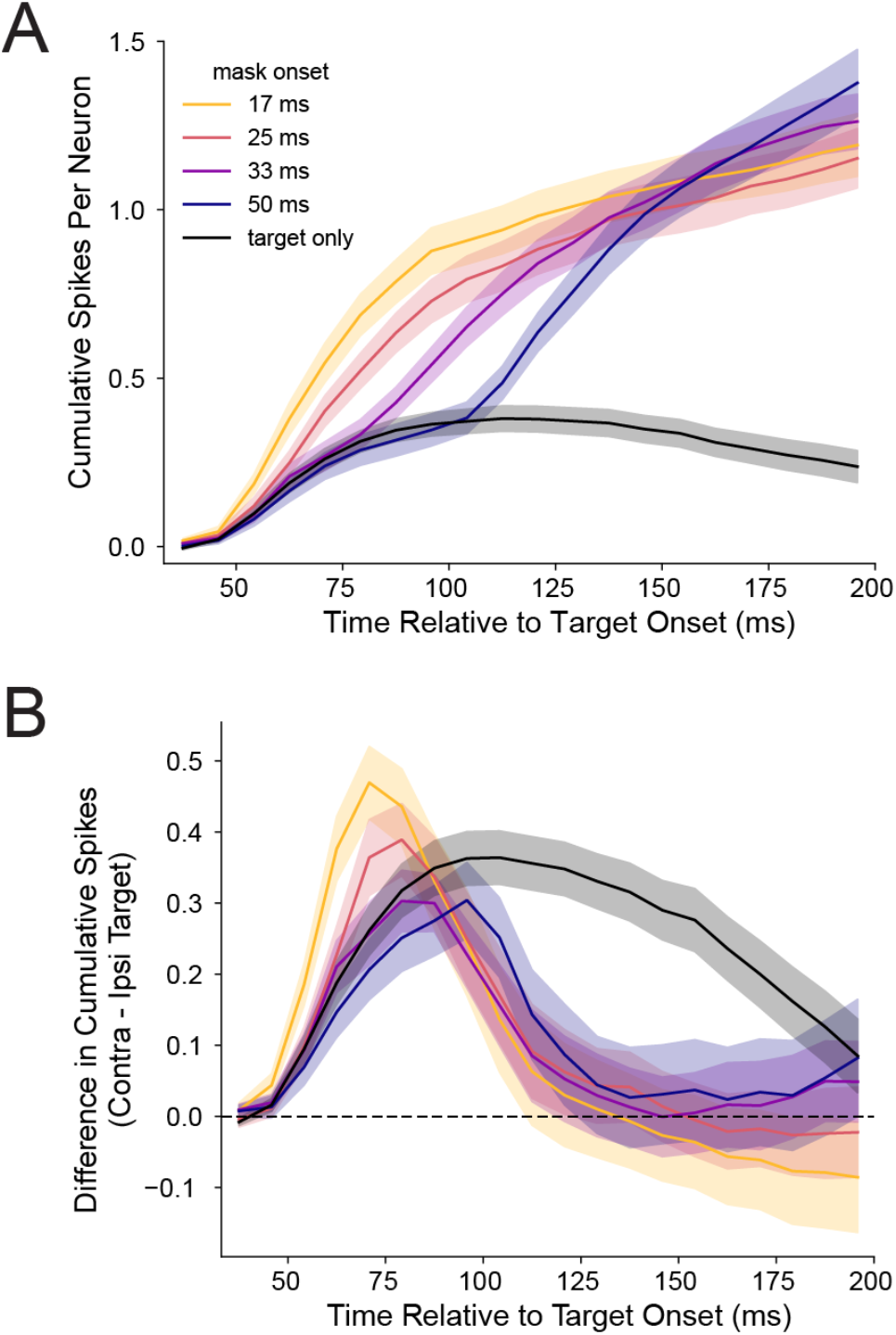
**(A)** Cumulative stimulus-evoked spikes over 200 ms per V1 neuron (n=58) following onset of a contralateral target. Spike number was calculated by integrating the trial-averaged post-stimulus time histogram after baseline subtraction for each neuron, and then averaging across neurons. Shaded region surrounding each line represents the standard error of the mean. (B) The difference in the value quantified in A was calculated between trials with a contralateral or ipsilateral target (see also Fig. 3B,C).

**Figure S7.**
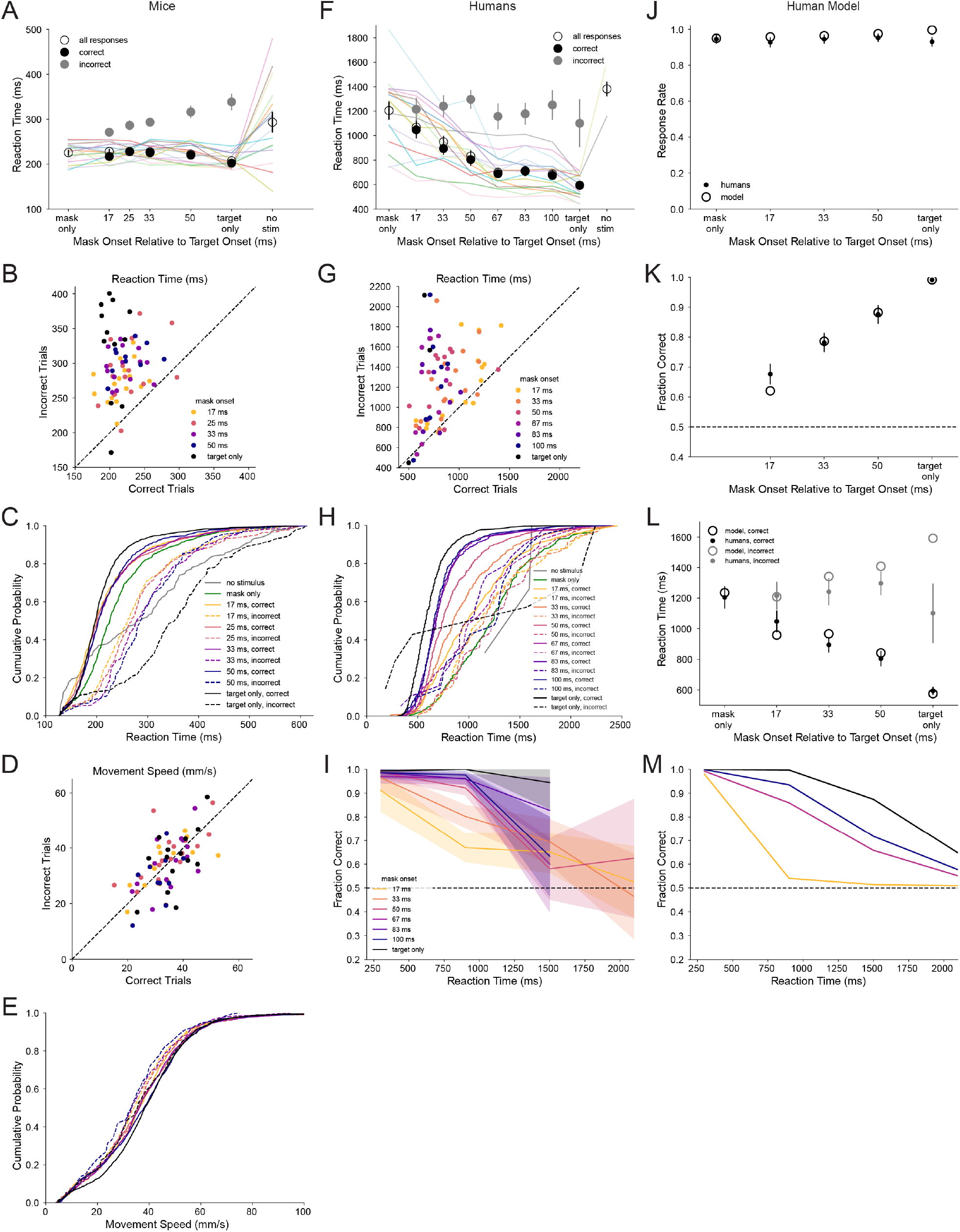
**(A)** Median reaction times for all responses, correct responses, and incorrect responses from the 16 mice used for Fig. 1B,C. Circles are means across mice and error bars represent standard error of the mean. **(B)** Comparison of median reaction times for correct and incorrect responses for each mouse. Each circle is data for one condition (mask onset or target only) from one mouse. **(C)** Cumulative probability distributions of reaction times pooled across the mice for correct (solid lines) and incorrect (dashed lines) responses. Reaction times on mask-only and catch trials are also shown. **(D,E)** Same as B,C for movement speed. **(F-H)** Same as A-C for the reaction times of 16 human subjects. **(I)** Fraction correct versus reaction time after pooling reaction times across all humans in 600 ms bins. Shaded region surrounding each line is the 95% confidence interval given the fraction correct values and the number of pooled trials for each bin (the median number of trials across mask onset conditions was 135, 296, 30, and 28 for the time bins from left to right after excluding conditions from a bin if there were less than 15 trials). **(J-L)** Response rate, accuracy, and reaction times of the dual accumulator model (open circles) shown in Fig. 3F fit to human behavior data (filled circles with error bars representing the standard error of the mean). **(M)** Accuracy versus reaction time for the model, corresponding to I for humans.

**Figure S8.**
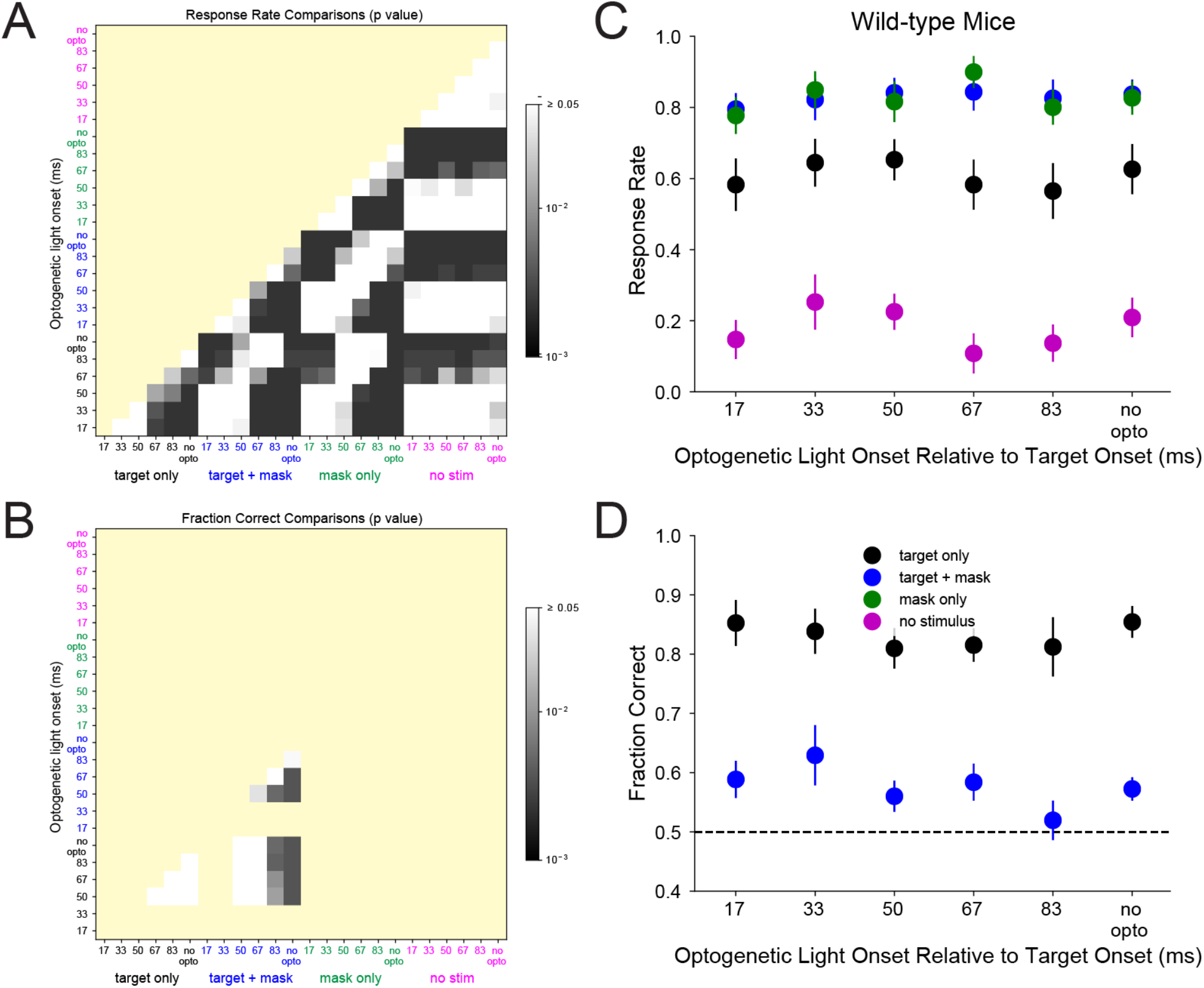
**(A,B)** p-values from comparison of response rates (A) and accuracy (B) for the conditions shown in Fig. 4A,B, using statistical procedures described in the legend for Fig. S2 A,B. **(C,D)** Effect of optogenetic light stimulus on response rate and accuracy in wild-type mice (n=8) for the same conditions shown in Fig. 4A,B.

**Table S1.**
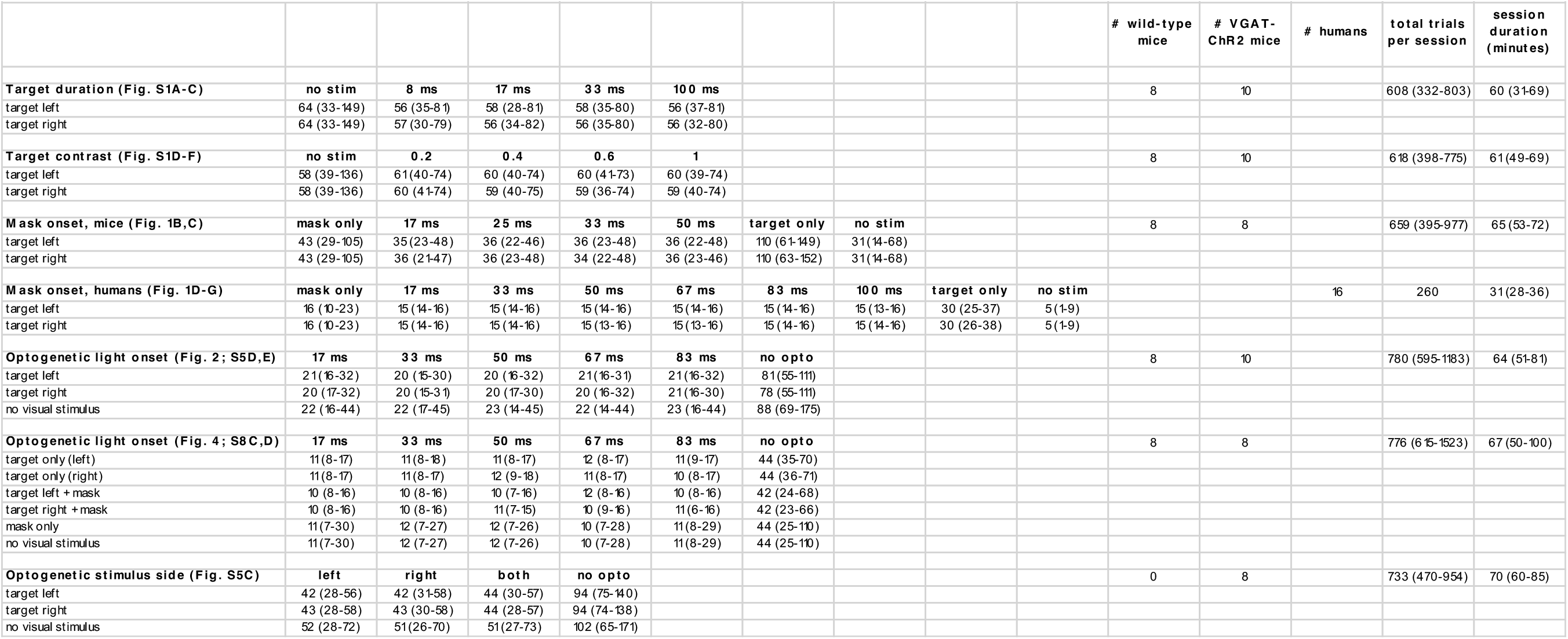
Median number of trials (and range) per session.

**Table S2.**
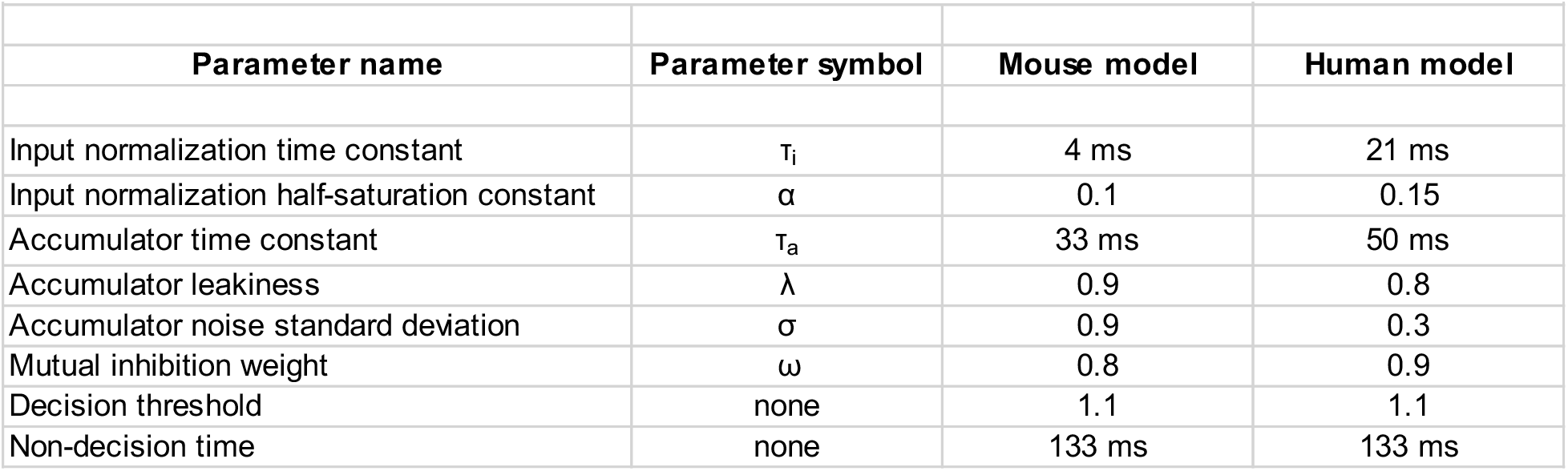
Values of fit parameters for the dual accumulator model.

## Notes

### Competing Interest Statement

The authors have declared no competing interest.

### Summary of Updates

New experiment added in which humans are tested on the same backward visual masking task previously used with mice.

